# Oral immunization with rVSV bivalent vaccine elicits protective immune responses, including ADCC, against both SARS-CoV-2 and Influenza A viruses

**DOI:** 10.1101/2023.07.14.549076

**Authors:** Maggie Jing Ouyang, Zhujun Ao, Titus A. Olukitibi, Peter Lawrynuik, Christopher Shieh, Sam Kung, Keith Fowke, Darwyn Kobasa, Xiaojian Yao

**Author notes:** To whom correspondence should be addressed: X-j. Yao, Laboratory of Molecular Human Retrovirology, Department of Medical Microbiology, Max Rady College of Medicine, Rady Faculty of Health Sciences, University of Manitoba.

## Abstract

COVID-19 and influenza both cause enormous disease burdens, and vaccines are the primary measures for their control. Since these viral diseases are transmitted through the mucosal surface of the respiratory tract, developing an effective and convenient mucosal vaccine should be a high priority. We previously reported a recombinant vesicular stomatitis virus (rVSV)-based bivalent vaccine (v-EM2/SPΔC1_Delta_) that protects animals from both SARS-CoV-2 and influenza viruses via intramuscular and intranasal immunization. Here, we further investigated the immune response induced by oral immunization with this vaccine and its protective efficacy in mice. The results demonstrated that the oral cavity delivery, like the intranasal route, elicited strong and protective systemic immune responses against SARS-CoV-2 and influenza A virus. This included high levels of neutralizing antibodies (NAbs) against SARS-CoV-2, as well as strong anti-SARS-CoV-2 spike protein (SP) antibody-dependent cellular cytotoxicity (ADCC) and anti-influenza M2 ADCC responses in mice sera. Furthermore, it provided efficient protection against challenge with influenza H1N1 virus in a mouse model, with a 100% survival rate and a significant low lung viral load of influenza virus. All these findings provide substantial evidence for the effectiveness of oral immunization with the rVSV bivalent vaccine.

## Introduction

The global battle against the COVID-19 pandemic has lasted for three years. Waves of emerging variants break through the protection obtained from previous vaccination or infection^1^. Although COVID-19 vaccines attenuate illness severity,^1, 2^ there are still significant numbers of new cases and deaths worldwide.^3^ Improving vaccine efficiency is a top priority for effective COVID-19 control. The limitations of current SARS-CoV-2 vaccines include a spike protein (SP or S) antigen that has become highly mutated, as well as an intramuscular immunization route, which lacks protection that is strong and specific to the respiratory tract.^4, 5^ It has been reported that following a booster of the Pfizer vaccine (i.m.), salivary mucosal immunity against the Wuhan strain relied on serum-exuded IgG, but not on locally produced secretory IgA, because the vaccine failed to activate an effective mucosal immunity.^5^ Influenza is another important respiratory infectious disease causing a high disease burden. Influenza control faces similar problems annually: unpredictable prevailing strains and limited protection from intramuscular vaccines.

The natural portal of entry for SARS-CoV-2 and influenza virus is the mucous membranes lining the respiratory tract, including the nose, mouth, trachea, and lungs ^6, 7^. Therefore, the mucosa is the first line of defense against viruses. Theoretically, mucosal vaccines administered via the respiratory tract may elicit more robust local protective immune responses in comparison to intramuscular vaccines. Previous mice and macaques studies have reported that intranasal COVID-19 vaccines induced robust mucosal and systemic immune responses, especially tissue-resident memory T and B cells.^8, 9^ The immunized animals were completely protected against lethal SARS-CoV-2 challenge. Moreover, viral replication was not detected in the airways and lungs of immunized animals following viral challenge.^8, 9^ Recently, four mucosal COVID-19 vaccines have been approved for human emergency use and about 20 mucosal vaccines have reached clinical trials in humans.^10–13^ The approved vaccines from China (CanSino Biologics), India (Bharat Biotech), Iran (Razi Vaccine & Serum Res) and Russia (Sputnik V) are adenovirus-vectored. The CanSino vaccine is an aerosolized mist inhaled orally through the nose and mouth.^14, 15^ The Bharat vaccine is administered via nose drops.^11^ The Iranian (Razi Vaccine & Serum Res) and Russian (Sputnik V) vaccines both are nasal sprays.^10, 11, 16^ Oral vaccines, as one type of mucosal vaccine, can be easily administered to the mouth or gut without any device (such as a sprayer for the intranasal vaccine). It also can successfully induce immune responses. For example, Vaxart’s oral tablet COVID-19 vaccine that targets the mucosal epithelium of the small bowel has been demonstrated to generate broad cross-reactive T cell and mucosal IgA responses in a phase I clinical trial.^12^

The mucosal delivery of recombinant vesicular stomatitis virus (rVSV) has been an interesting area of research. The rVSV vaccine vector is a promising platform with some favorable characteristics, including its non-pathogenic nature and low pre-existing immunity in humans.^17^ The replacement of VSV-glycoprotein (G) with a viral antigen from the virus of interest may further reduce its tropism. Importantly, the safety of the rVSV vector has been demonstrated by many animal studies and clinical trials for the rVSV-based Zaire Ebolavirus vaccine (VSV-ZEBOV-GP, ERVEBO), which was approved by the FDA for human use in 2019.^18–22^ However, the VSV-ZEBOV-GP vaccine was administered intramuscularly. Also, it was reported that when the wild type VSV is administered intranasally, its neurotoxicity is still a concern because it carries the potential for brain infection via the olfactory tract above the nose cavity.^23^ Oral delivery of rVSV may carry a smaller risk than the intranasal route. It is worth noting that oral delivery of the rVSV vaccine has been reported to induce protective immune responses against the Sin Nombre virus and SARS-CoV-2 infection.^24, 25^ The rVSV-Sin Nombre virus (SNV) vaccine was delivered to deer mice via oral gavages, in which the SNV glycoprotein (G) replaced the VSV-G to mediate vaccine entry into the mucosa.^24, 26^ The rVSV-SARS2(+G) vaccine targeted the oral cavity mucosa through the VSV-G proteins that were incorporated in the virion’s surface (trans-complemented) and were responsible for the target cell tropism.^25^ We recently have reported a replication-competent rVSV-based bivalent vaccine (v-EM2/SPΔC1_Delta_) that effectively protected animals against both SARS-CoV-2 and influenza A virus.^27^ This bivalent vaccine has a modified Ebola virus glycoprotein (GP) that mediates rVSV entry to cells, including macrophages and dendritic cells (DCs). Our vaccine has elicited robust adaptive immune responses via either intramuscular or intranasal vaccination in mice or hamsters.

Vaccine-induced virus-specific antibodies provide anti-viral protection through not only neutralization, but also extra-neutralizing functions, such as antibody-dependent cellular cytotoxicity (ADCC), antibody-dependent cellular phagocytosis (ADCP), and complement-dependent cytotoxicity (CDC).^28^ Among them, ADCC has been linked to the resolution of and protection against several viruses, including SARS-CoV-2^29–33^, influenza virus^34, 35^, HIV-1^36^, and Ebola virus.^37^ Yu et al. reported that COVID-19 patients who had recovered from severe disease had greater ADCC activity than patients who had succumbed to severe disease ^30^. Vigon et al. discovered that COVID-19 patients who required ICU assistance had significantly enhanced levels of memory B cells, plasmablasts, as well as neutralizing antibodies against SARS-CoV-2, but also impaired ADCC activity^29^. These findings strongly suggest that ADCC activity should be used to evaluate potential vaccines against SARS-CoV-2 in addition to NAbs.

In this study, we investigated the immune responses induced by oral immunization with a rVSV-based bivalent vaccine (v-EM2/SPΔC1_Delta_) in mice and compared it to intranasal immunization. We found that oral immunization elicited strong systematic immune responses against the Delta variant and influenza H1N1 virus, including neutralizing antibodies (NAbs) and antibody-dependent cellular cytotoxicity (ADCC) activity, which were similar to the efficacy of the intranasal immunization route. Further, oral immunization provided complete protection against influenza A H1N1 viral challenge in mice, like the intranasal route, with a 100% survival rate, as well as less weight loss, and significantly lower viral loads in the lungs. In addition, we discovered that the antibodies targeting SP and M2e were positively correlated with ADCC activity against SARS-CoV-2 and influenza.

## Results

### Oral immunization with v-EM2/SPΔC1_Delta_ elicited a robust humoral immune response and neutralization against SARS-CoV-2 in mice

Given that the oral delivery of several rVSV-based vaccines have been reported to induce protective immune responses in animals;^24, 25^ we want to extend the vaccination routes of our rVSV-based bivalent vaccine v-EM2/SPΔC1_Delta_^27^ to include oral immunization. To this end, we immunized female BALB/c mice with v-EM2/SPΔC1_Delta_ via oral cavity (oral, 1×10^6^ 50% Tissue Culture Infectious Dose, TCID_50_) or intranasal (i.n., 1×10^5^ TCID_50_) routes in three groups (n=5 each group) for prime/boost: 1) i.n./i.n.; 2) i.n./oral; and 3) oral/oral (Fig.1A and B). The mice in the control group were given PBS. The interval between prime and booster immunization was 2 weeks. The prime sera (2 weeks post Prime) and booster sera (3 weeks post booster) were collected. Their antibody levels were measured by ELISA (Fig.1 and Fig.2). As expected, intranasal vaccination induced robust anti-SARS-CoV-2 RBD (Fig.1C, E, and G) and anti-SARS-CoV-2 S2 (Fig.1D and F) humoral immune responses. Importantly, oral immunization elicited high levels of anti-RBD and anti-S2 IgG, similar to i.n. immunization (Fig.1C-F). The booster significantly enhanced antibody responses in all v-EM2/SPΔC1_Delta_ immunized groups (Fig.1C and D). Although the oral/oral group showed slightly lower average OD450 values in prime sera, no significant difference was found in the booster sera antibody titers between oral/oral and other immunized groups (Fig.1C-F). However, it is worth mentioning that the dose of oral immunization (10^6^ TCID_50_) was ten times that of i.n. immunization (10^5^ TCID_50_). Moreover, the anti-RBD IgA levels also exhibited high similarity among the three immunized groups (Fig.1G). These findings indicated that mucosal immunization with the v-EM2/SPΔC1_Delta_ vaccine through the oral cavity could effectively induce anti-SARS-CoV-2 SP antibody responses in mice. In addition, we have investigated the duration of antibodies in three i.n. immunized mice that were not challenged with the influenza virus. We found the anti-RBD IgG levels in these mice were maintained at a high peak from week 3 to week 5 after booster (Fig.2A), suggesting that the high levels of anti-SARS-Cov-2 antibodies in mice were sustained during the observed period.

**Figure 1.**
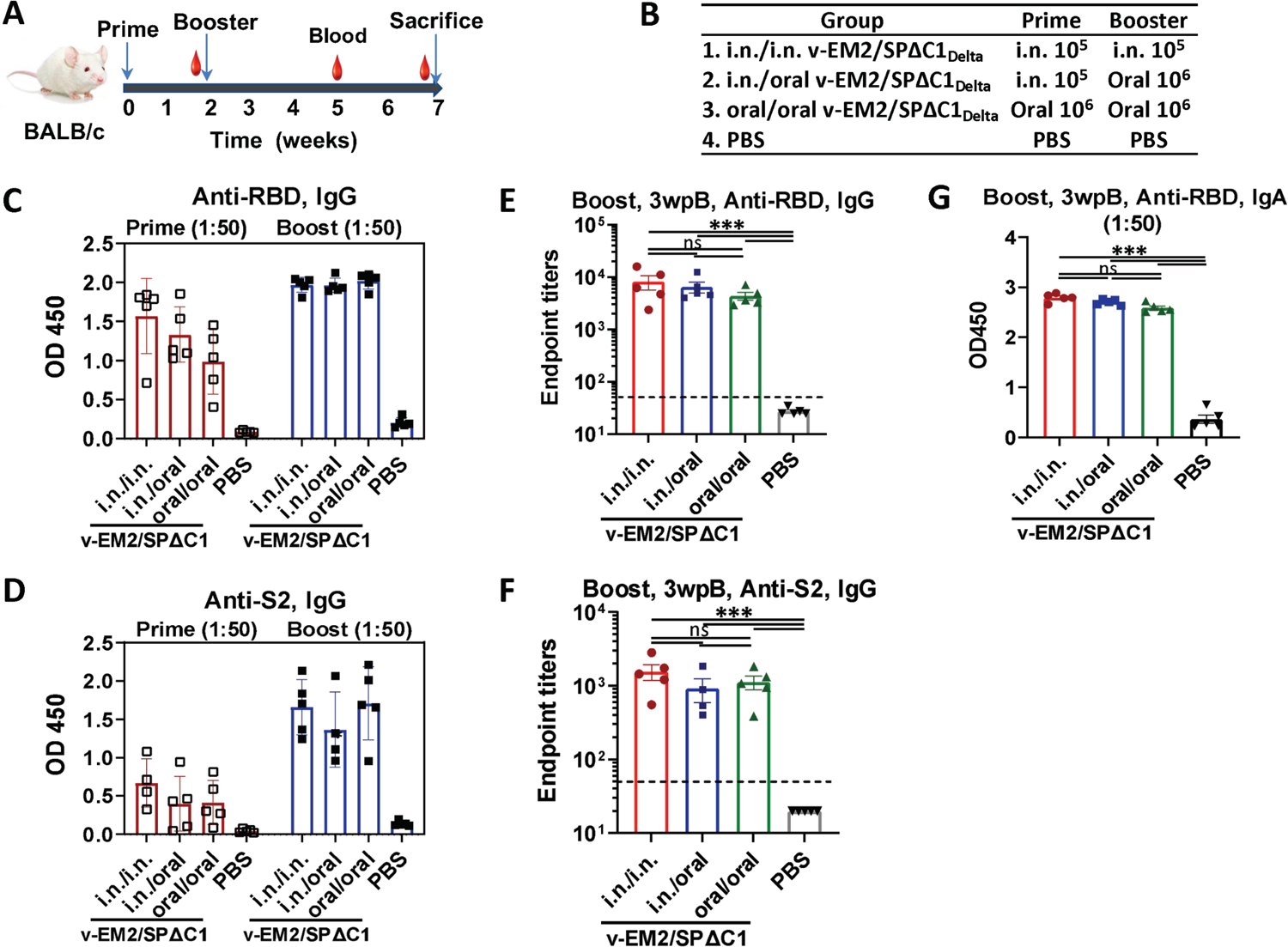
Oral immunization with vaccine candidate v-EM2/SPΔC1_Delta_ elicited robust anti-SARS-CoV-2 RBD and anti-S2 immune response in mice. A) Schematic of the immunization of bivalent rVSV vaccine candidate v-EM2/SPΔC1_Delta_ in mice. BALB/c mice (n=5-8 /group) were immunized with v-EM2/SPΔC1_Delta_ via oral cavity or intranasal routes at week 0 and 2 as indicated. The blood from mice in each group were collected on week 2 and 5, and the mice were sacrificed at week 7 (i.e. 5 weeks post-booster (5wpB)) and the blood was collected. B) The immunization groups are shown with the delivery route and vaccine dose. C-F) Sera after prime (week 2) and booster (week 5, i.e. 3 weeks post-booster (3wpB)) immunization were measured for anti-SARS-CoV-2 RBD IgG (C, E), IgA (G) levels and anti-S2 IgG (D, F) in OD450 or endpoint titers. Data represent mean ±SEM. Statistical significance was determined using one-way ANOVA test and Tukey’s test. *, P < 0.05; **, P < 0.01; ***, P < 0.001.

**Figure 2.**
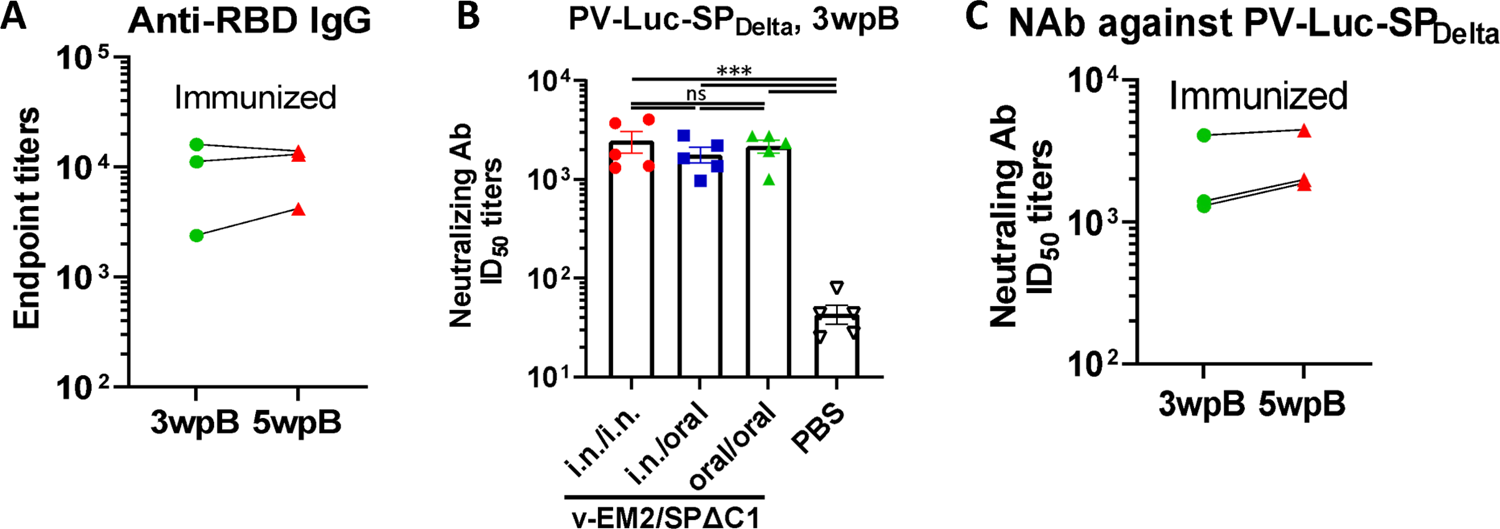
The duration and neutralization of antibodies against SARS-CoV-2 SP induced by oral immunization with v-EM2/SPΔC1_Delta_ in mice. **A)** The duration of anti-SARS-CoV-2 RBD (endpoint titers) in the i.n. immunized mice sera after booster immunization (3wpB and 5wpB) were measured. **B)** The neutralization titers (50% inhibitory dose, ID_50_) in immunized mice sera (3wpB) against pseudovirus PV-Luc-SP_Delta_ infection were determined. The serially diluted mouse sera were incubated with PV-Luc-SP_Delta_ (∼10^4^ RLU) and then, the mixtures (PV + Sera) were used to inoculate A549_ACE2_ cells. The infection of PV was determined by luciferase assay at 48∼66 hrs post-infection. The percentage of infection was calculated compared with no serum control. The 50% inhibition dose (ID_50_) neutralizing Ab titers were calculated by using sigmoid 4PL interpolation with GraphPad Prism 9.0, as described in Materials and Methods. **C)** The duration of neutralizing Ab ID_50_ titers in the i.n. immunized mice sera collected at 3wpB and 5wpB. Data represent mean ±SEM. Statistical significance was determined using one-way ANOVA test and Tukey’s test. *, P < 0.05; **, P < 0.01; ***, P < 0.001. wpB, week post Boost.

To evaluate if oral immunization with v-EM2/SPΔC1_Delta_ can induce protective immune responses, we measured neutralizing antibody (NAb) levels by using SP_Delta_-pseudotyped Luciferase+ HIV-based virus particles (PV-Luc-SP_Delta_) and human angiotensin-converting enzyme 2 (ACE2)-expressing cells A549_ACE2_^27, 38^. The results showed that all three immunized groups induced markedly high NAb titers against the Delta SP-peudotyped virus in comparison with the control group (Fig.2A). Further, the NAb levels of oral group mice had no significant difference from the other two groups (i.n./i.n. group and i.n./oral group). This finding confirmed that oral immunization with v-EM2/SPΔC1_Delta_ could effectively elicit protective NAbs against the SARS-CoV-2 Delta SP-peudotyped virus at a similar level to the i.n. vaccination. Like the RBD-binding antibodies (Fig. 2B), the NAb titers in the i.n. immunized mice sera showed stability from 3wpB to 5wpB (Fig. 2C), indicating that the high levels of neutralizing antibodies against Delta SP-pseudotyped virus in these immunized mice were sustained during the observed period.

### Oral immunization with v-EM2/SPΔC1_Delta_ induced ADCC activities against SARS-CoV-2 in mice

In addition to neutralization, other immune responses, such as ADCC that can kill virus-infected cells, have been demonstrated to contribute to vaccine protection.^39, 40^ Therefore, we have evaluated the ADCC activity induced by v-EM2/SPΔC1_Delta_ through a reporter system that used Jurkat-Lucia NFAT (nuclear factor of activated T cells)-CD16 (Invivogen) as effector cells. This T-cell reporter cell line, similar to natural killer (NK) cells, expresses the human Fc receptor FcγRIII (CD16; V158 allotype) on the cell surface, which can bind with the Fc region of human IgG and is cross-reactive with the Fc of mouse IgG^41^ (Fig.3A, upper panel). If the antibodies are bound with antigen-expressing target cells, the antigen-antibody-FcγR engagements will trigger the activation of NFAT pathway in Jurkat-Lucia cells, including the expression of NFAT-driven reporter luciferase.^30, 41, 42^ In this study, we used transient expression cells 293TN-SP_Delta_ (expressing the SPΔC of SARS-CoV-2 Delta variant) as target cells (Fig.3A, lower panel, lane 3). The serially diluted mice sera were first incubated with the target cells 293TN-SP_Delta_, and then the Jurkat-Lucia effector cells were added. ADCC responses were determined by measuring the induction of luciferase secreted from the activated Jurkat-Lucia cells in the supernatants, which were determined via relative light unit (RLU) changes compared to the no-serum control (only target cells and effector cells).

**Figure 3.**
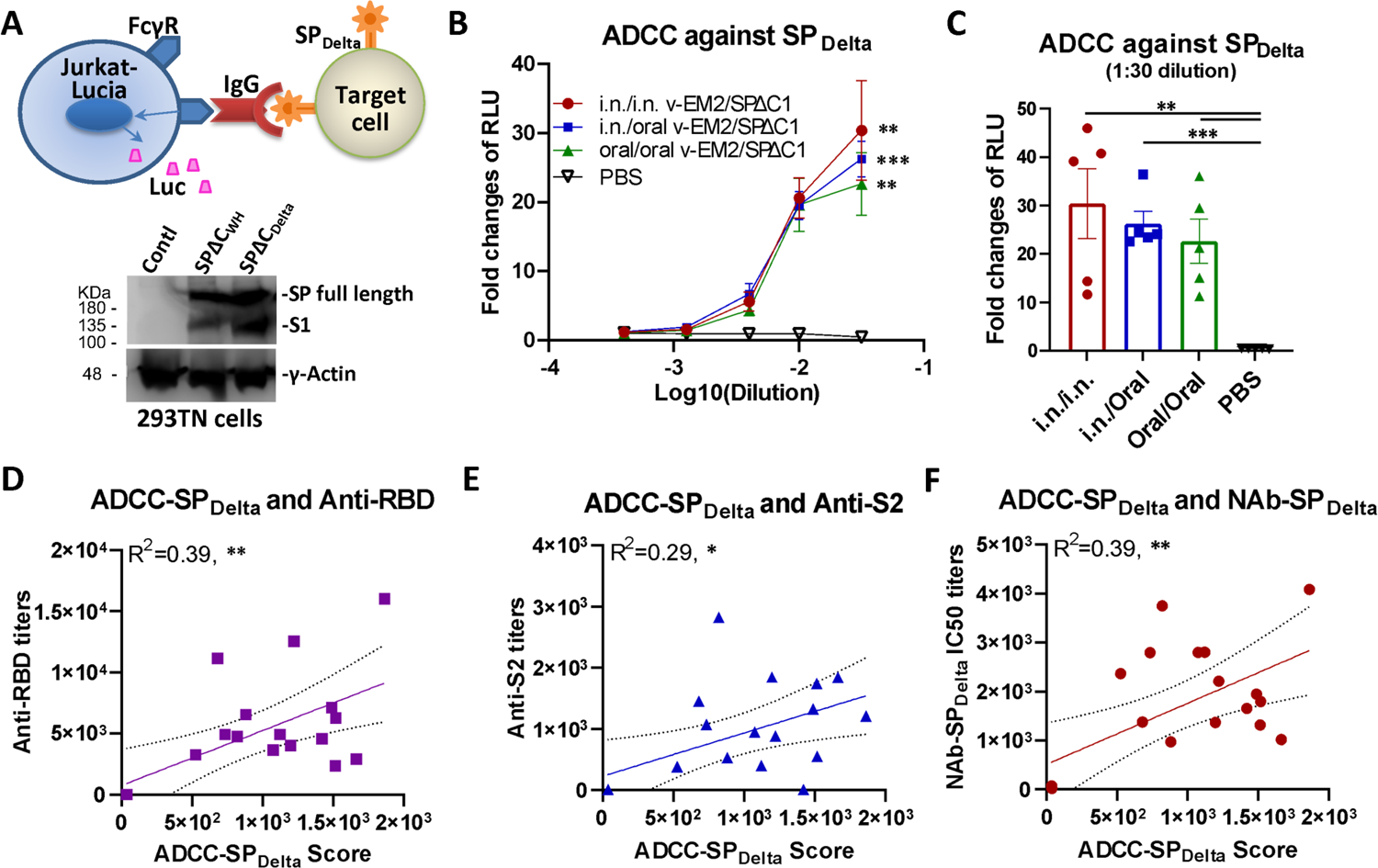
Oral immunization with v-EM2/SPΔC1_Delta_ induced the ADCC activities against SARS-CoV-2 SP_Delta_ in mice. **A)** The schematic diagram of Jurkat-Lucia cell-based ADCC activity reporter assay as described in Materials and Methods. The SP_Delta_-expressing cells are obtained from transfection and incubated with serially diluted mouse serum (containing SP-binding IgG). Jurkat-Lucia reporter cells expressing Fcγ receptor (FcγR) are added to the mixture of SP_Delta_-expressing cells and serum. The engagement of FcγR, IgG, and SP triggers the activation of Jurkat effector cells and the luciferase (Luc) expression. The ADCC activity is determined by measuring the secreted Luc in the cell culture supernatant (upper). The antigen SARS-CoV-2 spike proteins (SPΔC_WH_, SPΔC_Delta_) expression in target 293TN cells were determined by Western Blotting (lower panel) at 24 hrs post transfection using the anti-SARS-CoV-2 SP-NTD (Elabscience, Cat# E-AB-V1030). **B)** The ADCC activities against SARS-CoV-2 SP_Delta_ in the immunized mice sera (3wpB) were determined as described in (A). The ADCC induction (fold change) of sera from each group was calculated against the no-serum control. **C)** The ADCC against SP_Delta_ in individual mouse serum at 1:30 dilution. Data represents mean ± SEM. Statistical significance (B and C) was determined using an ordinary one-way (C) or two-way (B) ANOVA test and Tukey’s test. **D-F)** The ADCC activity against SP_Delta_ was positively correlated with the titers of anti-SP antibodies (RBD- or S2-binding) (D and E) or NAbs (F) of mice in all groups. The ADCC score was calculated as (fold-change × dilution factor). The ADCC score geomean of the first two dilutions was used. The correlation analysis was performed by Prism. The correlation coefficient (R^2^) and two-tailed P value were calculated via the Pearson method by Prism. *, P < 0.05; **, P < 0.01; ***, P < 0.001.

The results revealed that all immunized mice groups had significant ADCC activity against SP_Delta_ with a 23∼30 fold change at the first dilution (1:50, log_10_ dilution = −1.5), compared with the no-serum control (fold change of 1) (Fig.3B). Interestingly, although the oral/oral immunized mice sera showed a slightly lower ADCC activity than other i.n./i.n. and i.n./oral immunized groups, the differences were not statistically significant (Fig.3C), indicating a strong ADCC response was induced by oral immunization with v-EM2/SPΔC1_Delta_. In all, these results evidenced that oral vaccination with v-EM2/SPΔC1_Delta_ in mice elicited remarkable anti-SP_Delta_ antibody responses that not only have neutralizing activity, but also ADCC activity. Since the mucosa in the nasal cavity and oral cavity are so closely linked to each other, it is reasonable that they have the same efficacy in triggering host humoral immune responses. Our results indicated that these two immunization routes might be replaceable with each other or be used jointly in some scenarios.

ADCC is mediated by antigen-bound antibodies that can be recognized and captured by the Fc receptor on effector cells. It is reasonable to assume that only some of the vaccine-induced antibodies can trigger ADCC, so we wanted to know what kind of antibodies have a strong relationship with ADCC. To this end, we investigated the correlation between ADCC activity against SP_Delta_ and the titers of anti-SP antibodies (RBD- or S2-binding) (Fig.3D and E) or NAbs (Fig.3F) of mice in all groups. The results showed that ADCC activity against SP_Delta_ was positively correlated with anti-RBD antibodies (R^2^=0.39; **), anti-S2 antibodies (R^2^=0.29; *), and anti-SP_Delta_NAbs (R^2^=0.39; **) (Fig.3D-F). However, these correlations were not strong, suggesting that only a portion of these antibodies are related to ADCC. Similarly, the anti-SP_Delta_ NAbs only partially overlap with the ADCC-triggering antibodies. Our findings are consistent with previous reports that non-NAbs can also induce ADCC. ^30, 37^

### Oral immunization with v-EM2/SPΔC1_Delta_ effectively triggered humoral immune response and ADCC activity against the influenza virus

Given that the bivalent vaccine v-EM2/SPΔC1_Delta_ expresses both SARS-CoV-2 SPΔC1_Delta_ and four copies of the ectodomain of influenza virus M2 protein, we further verified the effectiveness of oral immunization against influenza by detecting anti-M2e IgG in the immunized mice sera (3wpB) by ELISA. The results showed that the oral/oral immunization route induced a high level of antibodies against M2e, although the level was slightly lower than that of the i.n./i.n. or i.n./oral immunization groups (Fig.4A).

**Figure 4.**
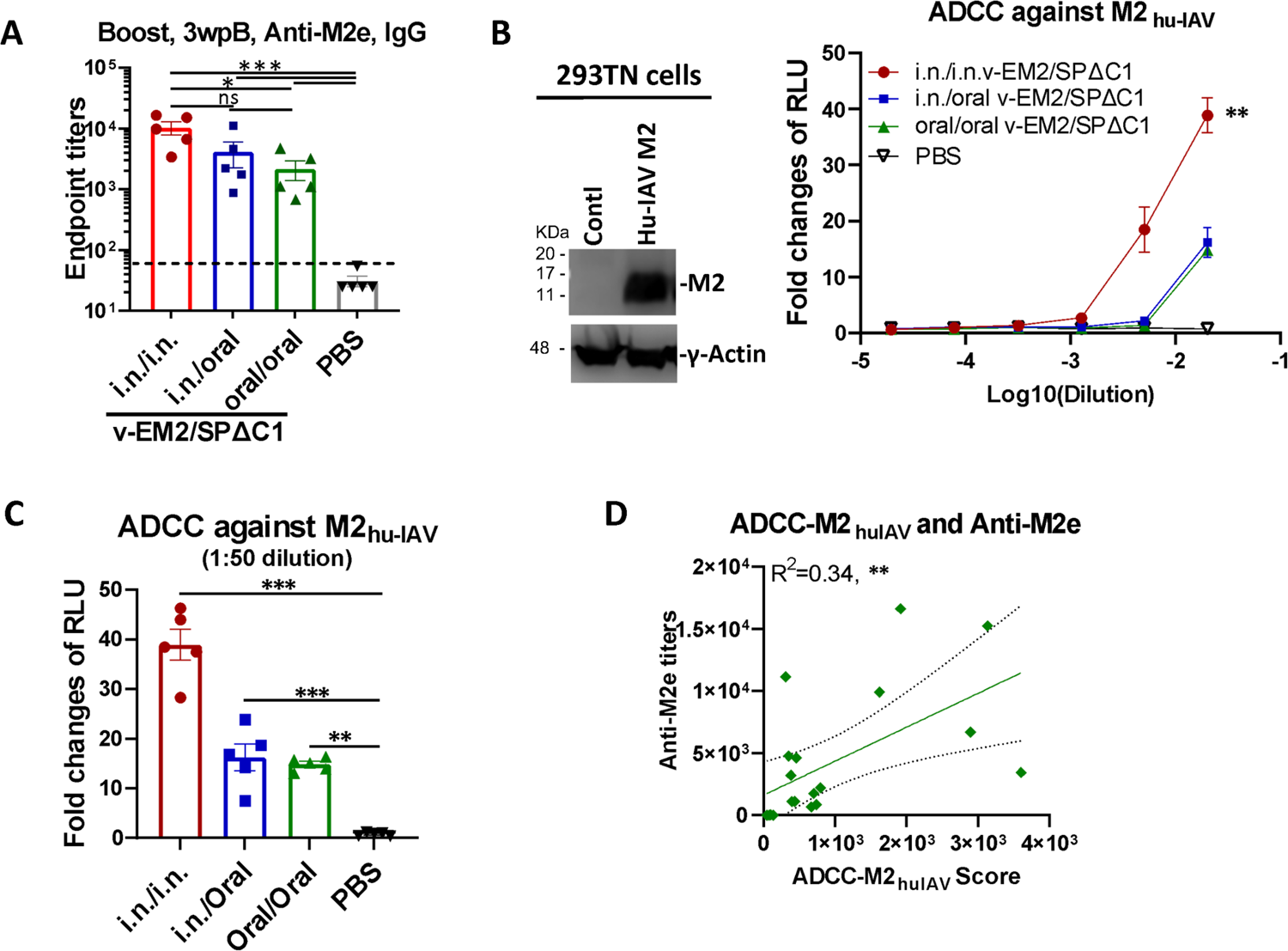
Oral immunization with vaccine candidate v-EM2/SPΔC1_Delta_ effectively triggered a humoral immune response against influenza virus in mice. **A)** The anti-influenza virus matrix-2 (M2) ectodomain (M2e) IgG levels in sera of the immunized mice at 3 weeks post booster (3wpB) as described in Fig.1A were measured by ELISA. **B)** The ADCC activities (fold change of RLU) against human influenza virus M2 (M2_hu-IAV_) in the immunized mice sera (3wpB) were determined as described in Fig.3A except using 293TN-M2 target cells. The human Influenza A virus (IAV) M2 protein expression in target cells was determined by Western Blotting using the sera of mouse immunized with v-EM2/ SPΔC1_Delta_ (containing anti-M2 antibodies) (left panel). The ADCC induction (fold change) of serum from each mouse was calculated against the no-serum control (right panel). **C)** The ADCC against M2_hu-IAV_ in individual mouse serum at 1:50 dilution. Data represents mean ± SEM. Statistical significance was determined using one-way (A, C) or two-way (B) ANOVA test and Tukey’s test. **D)** The correlation of ADCC activity against M2_hu-IAV_ and the titers of anti-M2e of mice in all groups. The ADCC score was calculated as described in Fig.3D. The correlation analysis was performed by Prism. The correlation coefficient (R^2^) and two-tailed P value were calculated via the Pearson method. *, P < 0.05; **, P < 0.01; ***, P < 0.001. wpB, week post Boost.

We then investigated ADCC activity against human influenza A virus M2 protein (M2_hu-IAV_) in the mice sera. Briefly, 293TN cells expressing Hu-IVA M2 (Fig.4B, left panel) were used as the target cells and incubated with serially diluted mice sera, followed by the addition of Jurkat-Lucia effector cells. ADCC activity was monitored by measuring the induction of luciferase secreted in the supernatants of the mix of 293TN-Hu-IAVM2/Jurkat-Lucia cell cultures (Fig 4B, right panel). The result disclosed that the i.n./i.n. immunized mice had a significantly high ADCC activity against M2_hu-IAV_ (fold change of 40 at 1:50 dilution). Further, the oral/oral and i.n./oral immunization sera both showed strong ADCC activity against M2_hu-IAV_ with a 15∼16 fold change at 1:50 dilution (Fig. 4C). Moreover, we found a positive correlation between the ADCC-M2_huIAV_ and the anti-M2e titers (R^2^=0.34, **) (Fig. 4D), indicating a part of anti-M2e antibodies contributed to the ADCC response as previously reported.^43–45^ These results also indicated that ADCC activity in the i.n./oral and oral/oral immunized mice play an important role for protection against influenza virus.

### Oral immunization with v-EM2/SPΔC1_Delta_ protected mice from lethal influenza challenge

To investigate the protective effects of oral immunization with v-EM2/SPΔC1_Delta,_ we performed a mouse challenge study by infecting immunized mice with a mouse-adapted influenza A virus PR8 (H1N1). Three different experimental groups of BALB/c mice were immunized with v-EM2/SPΔC1_Delta_ via i.n/i.n, oral/oral, or i.n/oral routes and a fourth control group was inoculated with PBS. The 4 groups were then infected with PR8 strain (3000 TCID_50_/mouse) intranasally on Day 42 (4 wpB) (Fig.5A). Weight loss and survival (Fig. 5B) was monitored every day for two weeks following the challenge. Excitingly, we found that the mice of three immunized groups all survived the challenge, indicating that either i.n. or oral immunization, or combined i.n./oral immunization, all effectively protected mice against influenza H1N1 virus infection. This was in contrast to the mice of the control group, which became moribund and reached the endpoint (weight loss of over 20%) within one week (Fig. 5B right panel). The immunized mice lost at most 8% of their initial weight 3 days post-infection (p.i.) and 4 days p.i., and then gradually regained weight to about 100% within two weeks. Notably, the oral/oral and i.n./oral groups did not show any less protection when compared with the i.n./i.n. group. This result is consistent with the above-described similarity between the anti-M2e antibody levels and the ADCC responses between the i.n. and the oral immunization routes (Fig. 4A and C).

**Figure 5.**
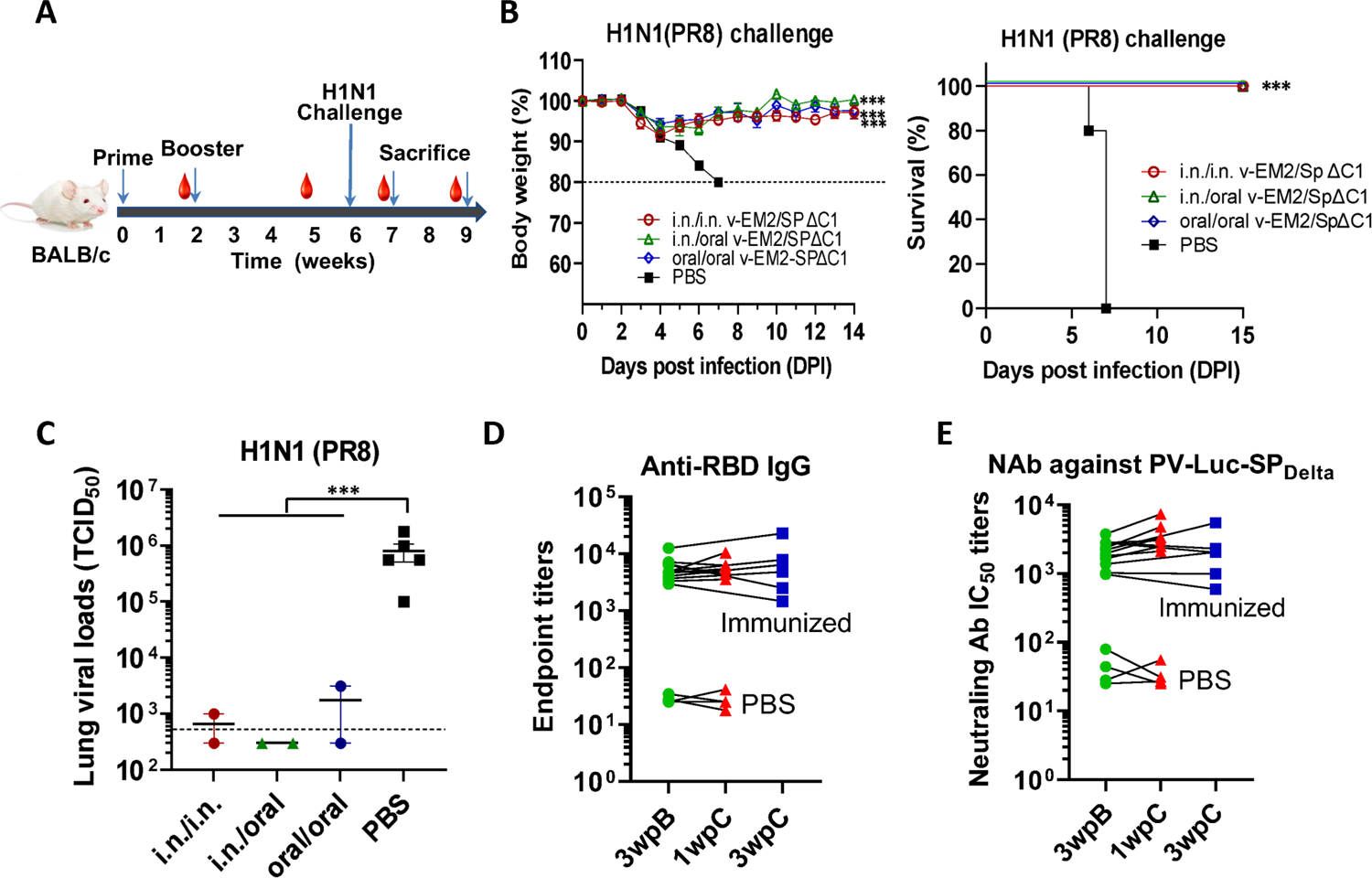
Oral immunization with v-EM2/SPΔC1_Delta_ protected mice from the lethal challenge of H1N1 influenza virus. **A)** Schematic of the immunization of bivalent rVSV vaccine v-EM2/SPΔC1_Delta_ and influenza challenge in mice. BALB/c mice (5/group) were immunized with v-EM2/SPΔC1_Delta_ via oral cavity or intranasal routes. At week 6, i.e. 4 weeks post-boost, mice of each group were intranasally infected with the H1N1 influenza virus PR8 (3×10^3^ TCID_50_). Then, the percentages of original body weight (**B**, left panel) and survival rates (**B**, right panel) of the mice were monitored daily for 2 weeks. **C)** Viral loads in the lung tissues of immunized mice (2/group) and PBS group (n=5) at 7 days post H1N1 challenge (1wpC) were determined with MDCK cells, as described in Materials and Methods. **(D** and **E)** The anti-RBD IgG endpoint titers (**D**) and NAb titers against PV-Luc-SP_Delta_ in mice sera before (3wpB) and after challenge (1wpC and 3wpC). The titers from the same mouse were linked with a line. Data shown is mean ± SEM. Statistical significance was determined using unpaired student T-test (C) or one-way ANOVA test and Tukey’s test (A and B). *, P < 0.05; **, P < 0.01; ***, P < 0.001. wpB, week post Boost. wpC, week post Challenge. TCID50, 50% Tissue Culture Infectious Dose.

The moribund mice in the control group were euthanized, and their blood and lungs were collected. Simultaneously, two mice from each immunized group were also euthanized at 7 days p.i. Their blood and lungs were collected as well. The lung viral loads of these infected mice were determined by TCID_50_ assay (Fig.5C). The results revealed that all the immunized mice had much lower viral loads in their lungs than the PBS control mice. This finding provides solid evidence that oral immunization with v-EM2/SPΔC1_Delta_ elicited protective immune responses in mice against influenza as strong as i.n. immunization.

In addition, we were curious whether infection with influenza virus could impact (either enhance or reduce) vaccine-induced immunity against SARS-CoV-2. To answer this question, we measured the anti-RBD IgG titers (Fig. 5D) and the NAb titers against SP_Delta_ (Fig. 5E) in the sera of mice challenged with influenza PR8 strain and tracked titer changes before (3wpB) and after challenge (1 and 3 weeks p.i). The results clearly showed that the levels of anti-RBD IgG and NAb against SP_Delta_ were similar or slightly higher at one week p.i. compared with those before the challenge. However, the anti-RBD IgG and NAb against SP_Delta_ levels returned back at 3wpi. Altogether, the above observations implied that the influenza virus infection did not significantly impact the vaccine-induced immunity against SARS-CoV-2.

## Discussion

The COVID-19 vaccines approved for emergency use in the last two years can only significantly reduce the severe illness and death, but not prevent the virus from spreading.^10, 11^ The major reason is breakthrough infections caused by immune-escaping SARS-CoV-2 variants. Another possible reason is the weak local immune response triggered by these intramuscularly administered vaccines.^5, 10, 11^ In addition, some evidence indicated that the release of circulating antibodies from the blood to the upper airway mucosa was limited by the restrictive blood-endothelial barrier in mice,^46^ suggesting the irreplaceable role of local immune response. Recently, mucosal vaccines have attracted more attention, because they efficiently prevent respiratory viral infections by improving the local immune response in addition to systemic responses.^12–15^ In this study, we demonstrated that robust systemic immune responses were induced by oral immunization with an rVSV-based bivalent vaccine (v-EM2/SPΔC1_Delta_) in mice against the SARS-CoV-2 Delta variant and H1N1 influenza A virus (IAV). Also, oral immunization with v-EM2/SPΔC1_Delta_ showed efficient protection against H1N1 influenza lethal challenge.

In detail, for the immune responses against SARS-CoV-2 Delta variants, highly similar profiles of anti-RBD and anti-S2 IgG/IgA titers (Fig.1E-G), the anti-SP_Delta_ NAb titers (Fig.2A) and anti-SP_Delta_ ADCC responses (Fig.3B and C) were observed between the oral (10^6^ TCID_50_/dose) and intranasal (10^5^ TCID_50_/dose) routes. Although we did not perform an authentic virus challenge experiment, there is solid evidence to support the efficacy of oral vaccination. Whether a smaller dose of oral immunization, like 10^5^ TCID_50_, can induce responses of a magnitude close to the i.n. group remains unknown. For the immune responses against influenza A virus, oral immunization achieved complete protection from a lethal H1N1 (PR8) challenge in mice (Fig.5A-C), high levels of anti-M2e antibodies in sera (Fig.4A), as well as strong anti-M2 ADCC activity (Fig.4B and C).

ADCC is another important anti-viral immune mechanism besides neutralization. Its protective contributions against SARS-CoV-2^29–33^ and influenza^34, 35^ have been reported. Through ADCC, Fc receptor (FcR)-bearing effector cells (Natural Killer cells, macrophages, and neutrophils) can recognize and kill target cells that are expressing viral antigens on their surface.^30, 47, 48^ The results provide evidence that oral immunization with our bivalent vaccine (v-EM2/SPΔC1_Delta_) in mice elicited a strong ADCC response against both the SARS-CoV-2 Delta variant (Fig.3B and C) and influenza virus (Fig. 4B and C), suggesting a major contribution of ADCC to protection. The initial step of ADCC is the engagement of Fc receptors on effector cells to specific antibodies that are already bound to the viral antigens on target cells. It is worth noting that the ADCC-inducing antibodies do not need to be neutralizing antibodies.^30, 37^ Our findings are in line with this: ADCC activity against SP_Delta_ showed a positive correlation with NAb (Fig.3F) but not in a perfect linear relationship, indicating the contribution of non-NAbs to ADCC. Another interesting discovery is that ADCC activity was positively correlated with both anti-RBD and anti-S2 antibodies (Fig.3D and E), implying the S1 (RBD) and S2 domains both can induce ADCC antibodies. This result suggests that the presence of both S1 (RBD) and S2 domains in a vaccine may be more beneficial than the single-domain vaccine in triggering ADCC-associated protection. To our surprise, the ADCC response against influenza-M2e elicited by the oral route was significantly lower than the intranasal group (Fig.4B and C). It appears to be correlated with the lower anti-M2e levels (Fig.4A). Given the complete protection achieved by both oral and i.n. immunization in the influenza virus challenge study (Fig. 5), it seems that such levels of peripheral anti-M2e and/or ADCC-triggering antibodies induced by oral vaccination, especially the local protective immune responses induced, (not reported here) are sufficient for protection. However, at this point, we have not investigated if oral immunization with the vaccine could provide equal protection from the SARS-CoV-2 infection, which is warranted for further study.

Our rVSV bivalent vaccine (v-EM2/SPΔC1_Delta_) is similar to the previously reported VSV-ZEBOV-GP (expressing Zaire Ebolavirus Glycoprotein/GP) vaccine that has been approved for clinical use.^18, 20–22^ We replaced the mucin-like domain of ZEBOV-GP with influenza matrix protein 2 ectodomain (M2e) and inserted the SARS-CoV-2 S in the upstream of VSV-L gene.^27^ Although Peng et al. recently reported poor immunogenicity for the oral delivery of the vaccine rVSV-SARS2, which expresses a single S protein,^25^ the difference is that our rVSV vaccine has an EBOV-GPΔM (deletion of the mucin-like domain) that was shown to be able to efficiently facilitate DC and macrophage targeting and induce more potent immune responses.^49, 50^ Therefore, it is still important to further investigate the mucosal immune response (such as the secreted IgA) and the memory T/B cells in the mucosa. Overall, this study showed successful oral immunization with the rVSV-based vaccine against both influenza and SARS-CoV-2 viral pathogens. In consideration of the efficacy, safety, and convenience of delivery, oral administration could be a promising option for rVSV-based vaccines, including v-EM2/SPΔC1_Delta_ vaccine.

In summary, we have demonstrated that oral immunization with the rVSV bivalent vaccine (v-EM2/SPΔC1_Delta_) in mice induced efficacious immune responses against both SARS-CoV-2 and influenza A virus, including high levels of antibodies that specifically bind to viral antigens and mediate neutralization and ADCC. Further we also clearly showed that vaccination via an oral route effectively protects mice from influenza virus challenge. As proof of concept, these findings provide evidence of good immunogenicity of the rVSV vaccine v-EM2/SPΔC1_Delta_ delivered in the oral cavity and high potential to induce protective immune responses.

### Limitations

Challenge in animals with authentic Delta variant was not included because of the limited resource of the level 4 containment lab. Mucosal immune responses and T/B cell immune responses were not included in this study, so local mucosal immunity induced by oral or nasal routes is still unknown. Our ADCC reporter assay could only can detect IgG-induced ADCC since the Jurkat effector cells express the CD16 (FCγRIII) receptor (IgG Fc receptor), so the data in this study only exhibited IgG-associated ADCC activity, which presented a part of the observed ADCC response, but not the IgA or IgE associated ADCC elements.

## Materials and methods

### Ethics statement

The animal experiments described were carried out according to protocols approved by the Central Animal Care Facility, University of Manitoba (Protocol Approval No. 20-034) following the guidelines provided by the Canadian Council on Animal Care. All animals were acclimated for at least one week before experimental manipulations and maintained with food and water ad libitum in a specific pathogen-free animal facility.

### Cells, plasmids, antibodies, recombinant proteins, and viruses

The HEK293T cells, human lung type II pulmonary epithelial A549_ACE2_ cell line,^38^ VeroE6, and MDCK cell lines were cultured in DMEM medium. Jurkat-Lucia NFAT-CD16 cells (Invivogen, Cat# jktl-nfat-cd16) were cultured in RPMI-1640 medium. The plasmids pCAGGS expressing SARS-CoV-2 SPΔC (deleted C-terminal 17 aa) from the original strain Wu-Han-1 (SPΔC_WH_), Delta variant B.1.617.2 (SPΔC_Delta_) and human influenza A virus M2 were constructed previously^27, 38, 51^. The antibodies used in this study included rabbit polyclonal antibody against SARS-CoV-2 SP/RBD (Sino Biological, Cat# 40150-R007), mouse monoclonal anti-SARS-CoV-2 S2 antibody (Abcam, Cat# 1A9: ab273433), human SARS-CoV-2 SP-NTD antibody (Elabscience, Cat# E-AB-V1030), mouse monoclonal anti-influenza A virus M2 antibody (Santa Cruz Biotech, Cat# 14C2: sc-32238), and anti-mouse-HRP antibody (GE Healthcare, Cat#NA931). Recombinant proteins included RBD peptide (RayBiotech, Cat# 230-30162), S1 subunit (RayBiotech, Cat# 230-01101), and S2 subunit (RayBiotech, Cat# 230-01103). Influenza M2e conserved peptide [M2e from human IAV (two copies), avian IAV (one copy), and swine IAV (one copy); 92 aa] was synthesized as previously described.^27, 52, 53^ The mouse-adapted influenza A/Puerto Rico/8/34 (H1N1) strain used for the animal challenge study was generated by reverse genetics as previously described^54^. The replication-competent recombinant vesicular stomatitis virus (rVSVs)-based COVID-19 vaccine v-EM2e/SPΔC1 was generated by inserting SPΔC_Delta_-I742A gene into the rVSV-EΔM-M2e vector as described previously.^27, 53^

### Mouse immunization with rVSV vaccine v-EM2/SPΔC1_Delta_

Oral immunization through the mouse oral cavity was performed by dropping the vaccine (25 µL/each) in the mouth under the tongue and into the cheek pouches after isoflurane anesthesia and keeping the mouse lying on its side for about 1 min until it woke up.^25^ Female BALB/c mice aged 6–8 weeks (five to eight per group) were immunized with bivalent rVSV vaccine v-EM2/SPΔC1_Delta_ on Day 0 and Day 14 via three different routes: orally (oral/oral, 1×10^6^ TCID_50_), intranasally (i.n./i.n., 1×10^5^ TCID_50_), and combining intranasally at prime and orally at boost (i.n./oral). Immunization with PBS was used as a placebo control. The blood of immunized mice was collected on Day 13 (prime), Day 35 (i.e., 3 weeks post-boost /3 wpB), and Day 49 (5 wpB). These sera were used to measure the RBD-, S2- and M2e-binding antibodies, the neutralizing antibodies, and antibody-dependent cellular cytotoxicity (ADCC).

### Measurement of vaccine-induced RBD-, S2- and M2e-binding antibodies titers

To determine RBD-, S2- and M2e-specific antibodies in mice sera, the sera of immunized mice were 3× serially diluted in primary antibody diluent (complete RPMI 1640 media with 0.2% (v/v) Tween 20). Enzyme-linked immunosorbent assay (ELISA) 96-well plates (NUNC Maxisorp, Thermo Scientific) were coated with recombinant proteins RBD, S2, or M2e (0.75 µg/mL) in coupling buffer (pH9.6, 50mM sodium carbonate-bicarbonate) at 4 °C overnight.^53^ After blocking at 37 °C for 2 hrs, the ELISA plates were washed and incubated with the diluted sera at 37 °C for 1 hr. Anti-mouse IgG-HRP antibodies (GE Healthcare, Cat#NA931; 1:5000) were used to detect the antibodies binding to RBD, S2, or M2e. After incubation with the substrate tetramethylbenzidine (TMB) solution (Mandel Scientific) and termination with stop solution, the absorbance at 450 nm (OD450) of each well was measured. IgA antibody levels were determined by using the mouse IgA ELISA kit (Thermofisher, Cat#88-50450-88). The endpoint titers of mouse sera were calculated using the interpolation in GraphPad Prism 9.0 with a cutoff of 2.5 times the Mean-negative.

### SP pseudovirus production and titration

The SARS-CoV-2 SP-pseudotyped HIV-based luciferase-expressing (Luc) pseudovirus (PV-Luc-SP_Delta_) was produced by co-transfecting 293T cells in a 6-well plate with the pCAGGS plasmid expressing SPΔC protein from B.1.617.2 (Delta)^55^ (0.5 µg/well) and a Luc-expressing HIV vector (pNL4-3-R-E-Luc)^56^ (1.0 µg/well) as described previously.^27^ The supernatants containing SP-pseudovirus were harvested at 72 hrs post-transfection, filtered (0.45 μm filter), aliquoted, and stored at −80 °C. The pseudovirus was titrated on A549_ACE2_ cells by a modified method.^27, 57, 58^ Briefly, the pseudovirus was 2× serially diluted in 25 μL of complete DMEM and mixed with A549_ACE2_ cells (1.25×10^4^/well; 50 μL) and polybrene (5 µg/mL) in a 96-well plate for transduction. After overnight incubation, cells were fed with fresh complete DMEM. At 48∼66 hrs post-infection, cells were lysed in luciferase lysis buffer (Promega; 30 μL /well), and the luciferase relative light unit (RLU) of the lysates was measured using the luciferase assay system (Promega) and Polerstar optima microplate reader (BMG BioLabtech) according to the manufacturer’s instructions. The HIV-1 p24 in PVs was also quantified by ELISA as previously reported.^38, 59^

### Pseudovirus-based neutralization assays against SARS-CoV-2

The neutralization assay was performed based on SARS-CoV-2 SP-pseudotyped HIV-Luc pseudovirus and A549_ACE2_ cells according to the previous method.^27, 57, 58^ Briefly, 2× serially diluted inactivated mouse sera (starting from 1:50 dilution, 25 µL) was pre-incubated with PV-Luc-SPΔC (about 10^4^ RLUs, 25 µL) in complete DMEM with polybrene (5 µg/mL) in a 96-well plate at 37°C for 1.5 h, then A549_ACE2_ (1.25×10^4^ cells/well, 50 µL) was added to each well and incubated at 37 °C for 48 hrs. The luciferase RLU in cells was measured by using Luciferase Assay System (Promega). The neutralizing titers or half-maximal inhibitory dilution (ID_50_) were defined as the reciprocal of the serum maximum dilution that reduced RLUs by 50% compared with no-serum (virus and cell) controls. The ID_50_ was calculated by using sigmoid 4PL interpolation with GraphPad Prism 9.0.

### Antibody-dependent cellular cytotoxicity (ADCC) reporter assay

The ADCC reporter assay was performed by using Jurkat-Lucia NFAT-CD16 cells (human FcγRIII, V158) (Invivogen, Cat# jktl-nfat-cd16) as effector cells according to the manufacturer’s instruction with modifications. The engagements among Jurkat-Lucia FcγRIII (CD16), antibodies (serum), and antigen-expressing target cells activate luciferase expression driven by NFAT (nuclear factor of activated T cells) in Jurkat-Lucia.^30, 41, 42^ In this study, we make use of the ADCC reporter system based on the cross-reactivities of mouse IgG to human FcγRs.^41^ Briefly, one day before the assay, 293TN cells (5×10^5^/well, 6-well plate) were transfected with each plasmid expressing SPΔC of SARS-CoV-2 Delta variant (1.2 µg DNA) or human influenza A virus M2 (5 µg DNA) using Lipofectamine 3000 (Invitrogen, USA) to produce 293TN-S target cells. S expression was semi-quantified by a western blot the next day (Fig. 3A and 4B). The mouse sera were 3× serially diluted in complete DMEM (start from 1:30, 50 µL/well) in a 96-well plate, and then mixed with target cells 293TN-S or 293TN-M2 (5×10^4^, 50 µL/well), incubated at 37 °C for 1 hr. The Jurkat-Lucia effector cells (1.5×10^5^, 50 µL/well) were added and cultured for additional 16 hrs at 37 °C with 5% CO2. ADCC activity was determined by measuring the secreted luciferase from activated Jurkat-Lucia cells in the supernatant (30 µL/well). The relative light unit (RLU) was detected by Quanti-Luc substrate solution (25 µL/well, Invivogen) using a Polerstar optima microplate reader (BMG BioLabtech). The background well (only medium) reading was deducted. The no-serum controls were wells containing target cells and Jurkat-Lucia cells. The fold of change (ADCC induction) was calculated as RLU (test − background)/RLU (no-serum control − background). The ADCC activity of sera was calculated as the geometric mean of the ADCC score (fold change × dilution factor) from two dilutions.

### Mouse challenge with influenza H1N1 virus PR8

In the influenza virus challenge study, the four groups of female mice (5 for each group) that were immunized with v-EM2/SPΔC1 or PBS as described above were intranasally infected with a mouse-adapted strain A/Puerto Rico/8/34 (PR8, H1N1) (3×10^3^ PFU/mouse) on Day 42 (4 wpB). The weight and survival of mice were monitored daily for 2 weeks after the challenge. The mice from the PBS group reached the endpoint (moribund or weight loss over 20%) at 6∼7 days post-challenge (1week post-challenge, 1wpC), and then were euthanized with two mice of each vaccinated group. The blood/sera and lungs were collected and stored at −80°C for viral load assay, ELISA, and other assays.

### Measurement of viral load in the mouse lungs (TCID_50_)

To determine the viral load of influenza H1N1 (PR8) in mice, we homogenized each mouse lung in ice-cold 1 ml of DMEM using a tenbroeck glass homogenizer. After spin, the supernatants were 10× serially diluted (starting from 1:10^2^) in the influenza virus infection media (DMEM, 1%P/S, 1 µg/mL TPCK-trypsin). The inoculums (100 µL/well) were added into the 96-well plate with 95% confluent MDCK cells and incubated at 37 °C. After 3 days, plates were fixed, stained with 2% crystal violet, and scored for cytopathic effect (CPE). The 50% Tissue Culture Infectious Dose (TCID_50_) titers were calculated using a modified Reed and Muench method.^60, 61^

### Statistical analysis

Statistical analysis of antibody/cytokine levels was performed using the unpaired Student T-test for two groups comparison (P ≥ 0.05 considered significant) by GraphPad Prism software. The statistical analysis for the endpoint titers, neutralizing antibodies, and mouse weight loss was performed using the one-way ANOVA multiple comparison test followed by Tukey’s test with Prism. For the ADCC responses, the two-way ANOVA multiple comparison test followed by Tukey’s test was used. For the survival rate, the Log-rank (Mantel-Cox) test was used. The correlation coefficient (r and R^2^) and two-tailed P value were calculated via the Pearson method with Prism.

## Data Availability

The data sets generated and/or analyzed during the current study are available from the corresponding author.

## Acknowledgments

This work was supported by the Canadian 2019 Novel Coronavirus (COVID-19) Rapid Research Funding (OV5-170710) by the Canadian Institutes of Health Research (CIHR, cihrirsc.gc.ca/e/193.html), Research Manitoba (researchmanitoba.ca), and CIHR COVID-19 Variant Supplement grant (VS1-175520) (to X.Y.). This work was also supported by the Manitoba Research Chair Award from Research Manitoba (RM) to X.Y.. T.O. is a recipient of the University Manitoba Graduate scholarship.

## Author contributions

Conceptualization, X.Y., M.O., S.F., K.F., and D.K.; The rVSV vaccine preparations were performed by Z.A., and X.Y.; Animal immunization and challenge studies, M.O., and T.O.; Neutralization, and ADCC experiments, M.O., T.O., and Z.A.; manuscript writing—review and editing, M.O., Z.A., T.O., P.L., C.S., S.F., K.F., D.K., and X.Y., Funding acquisition, X.Y., S.F., K.F., and D.K.; All authors have read and agreed to the published version of the manuscript.

## References

1. Andrews, N. et al. Covid-19 Vaccine Effectiveness against the Omicron (B.1.1.529) Variant. New England Journal of Medicine 386, 1532–1546 (2022).

2. Buchan, S.A. et al. Estimated Effectiveness of COVID-19 Vaccines Against Omicron or Delta Symptomatic Infection and Severe Outcomes. JAMA Network Open 5, e2232760–e2232760 (2022).

3. Organization, W.H. WHO Coronavirus (COVID-19) Dashboard. https://covid19.who.int/table (2022).

4. Tang, J. et al. Respiratory mucosal immunity against SARS-CoV-2 after mRNA vaccination. Science Immunology 7, eadd4853 (2022).

5. Azzi, L. et al. Mucosal immune response after the booster dose of the BNT162b2 COVID-19 vaccine. eBioMedicine 88 (2023).

6. Huang, N. et al. SARS-CoV-2 infection of the oral cavity and saliva. Nat Med 27, 892–903 (2021).

7. Killingley, B. & Nguyen-Van-Tam, J. Routes of influenza transmission. Influenza Other Respir Viruses 7 **Suppl 2**, 42–51 (2013).

8. Mao, T., et al. Unadjuvanted intranasal spike vaccine booster elicits robust protective mucosal immunity against sarbecoviruses. *bioRxiv*, 2022.2001.2024.477597 (2022).

9. Le Nouën, C., et al. Intranasal pediatric parainfluenza virus-vectored SARS-CoV-2 vaccine candidate is protective in macaques. *bioRxiv*, 2022.2005.2021.492923 (2022).

10. Waltz, E. China and India approve nasal COVID vaccines — are they a game changer? Nature 609, 450 (2022).

11. Waltz, E. How nasal-spray vaccines could change the pandemic. Nature 609, 240–242 (2022).

12. Johnson, S., et al. SARS-CoV-2 oral tablet vaccination induces neutralizing mucosal IgA in a phase 1 open label trial. *medRxiv*, 2022.2007.2016.22277601 (2022).

13. Kar, S. et al. Oral and intranasal vaccines against SARS-CoV-2: Current progress, prospects, advantages, and challenges. Immunity, inflammation and disease 10, e604 (2022).

14. Li, J.-X. et al. Safety and immunogenicity of heterologous boost immunisation with an orally administered aerosolised Ad5-nCoV after two-dose priming with an inactivated SARS-CoV-2 vaccine in Chinese adults: a randomised, open-label, single-centre trial. The Lancet Respiratory Medicine 10, 739–748 (2022).

15. Jin, L., et al. Antibody Persistence and Safety through 6 Months after Heterologous Orally Aerosolised Ad5-nCoV in individuals primed with two-dose CoronaVac previously. *medRxiv*, 2022.2007.2026.22278072 (2022).

16. Banihashemi, S.R. et al. Safety and Efficacy of Combined Intramuscular/Intranasal RAZI-COV PARS Vaccine Candidate Against SARS-CoV-2: A Preclinical Study in Several Animal Models. Front Immunol 13, 836745 (2022).

17. Fathi, A., Dahlke, C. & Addo, M.M. Recombinant vesicular stomatitis virus vector vaccines for WHO blueprint priority pathogens. Human vaccines & immunotherapeutics 15, 2269–2285 (2019).

18. Suder, E., Furuyama, W., Feldmann, H., Marzi, A. & de Wit, E. The vesicular stomatitis virus-based Ebola virus vaccine: From concept to clinical trials. Human vaccines & immunotherapeutics 14, 2107–2113 (2018).

19. Yon C. Yu; Pierre E. Rollin; Brian H. Harcourt; Robert L. Atmar; Beth P. Bell; Rita Helfand; Inger K. Damon; Sharon E. Frey., M.J.C.C.M.C.A.N.W.J.W.D.A.J.R.L.M.D.C.-O.M.P.E.E. Use of Ebola Vaccine: Recommendations of the Advisory Committee on Immunization Practices, United States, 2020. MMWR Recomm Rep 70, 1–12 (2021).

20. Pinski, A.N. & Messaoudi, I. Therapeutic vaccination strategies against EBOV by rVSV-EBOV-GP: the role of innate immunity. Current Opinion in Virology 51, 179–189 (2021).

21. Jones, S.M. et al. Assessment of a Vesicular Stomatitis Virus-Based Vaccine by Use of the Mouse Model of Ebola Virus Hemorrhagic Fever. The Journal of Infectious Diseases 196, S404–S412 (2007).

22. Geisbert, T.W. et al. Vesicular Stomatitis Virus-Based Ebola Vaccine Is Well-Tolerated and Protects Immunocompromised Nonhuman Primates. PLOS Pathogens 4, e1000225 (2008).

23. Durrant, D.M., Ghosh, S. & Klein, R.S. The Olfactory Bulb: An Immunosensory Effector Organ during Neurotropic Viral Infections. ACS chemical neuroscience 7, 464–469 (2016).

24. Warner, B.M. et al. Oral Vaccination With Recombinant Vesicular Stomatitis Virus Expressing Sin Nombre Virus Glycoprotein Prevents Sin Nombre Virus Transmission in Deer Mice. Frontiers in Cellular and Infection Microbiology 10 (2020).

25. Peng, K.W. et al. Boosting of SARS-CoV-2 immunity in nonhuman primates using an oral rhabdoviral vaccine. Vaccine 40, 2342–2351 (2022).

26. Kleinfelter, L.M. et al. Haploid Genetic Screen Reveals a Profound and Direct Dependence on Cholesterol for Hantavirus Membrane Fusion. mBio 6, e00801–00815 (2015).

27. Ao, Z. et al. A Recombinant VSV-Based Bivalent Vaccine Effectively Protects against Both SARS-CoV-2 and Influenza A Virus Infection. Journal of virology 96, e0133722 (2022).

28. Zohar, T. & Alter, G. Dissecting antibody-mediated protection against SARS-CoV-2. Nature reviews. Immunology 20, 392–394 (2020).

29. Vigón, L. et al. Impaired Antibody-Dependent Cellular Cytotoxicity in a Spanish Cohort of Patients With COVID-19 Admitted to the ICU. Frontiers in Immunology 12 (2021).

30. Yu, Y. et al. Antibody-dependent cellular cytotoxicity response to SARS-CoV-2 in COVID-19 patients. Signal Transduction and Targeted Therapy 6, 346 (2021).

31. Tso, F.Y. et al. Presence of antibody-dependent cellular cytotoxicity (ADCC) against SARS-CoV-2 in COVID-19 plasma. PLoS One 16, e0247640 (2021).

32. Hagemann, K. et al. Natural killer cell-mediated ADCC in SARS-CoV-2-infected individuals and vaccine recipients. European journal of immunology 52, 1297–1307 (2022).

33. Cui, T. et al. Potential of Antibody-Dependent Cellular Cytotoxicity in Acute and Recovery Phases of SARS-CoV-2 Infection. Infectious Diseases & Immunity 2, 74–82 (2022).

34. Gao, R., Sheng, Z., Sreenivasan, C.C., Wang, D. & Li, F. Influenza A Virus Antibodies with Antibody-Dependent Cellular Cytotoxicity Function. Viruses 12 (2020).

35. Von Holle, T.A. & Moody, M.A. Influenza and Antibody-Dependent Cellular Cytotoxicity. Front Immunol 10, 1457 (2019).

36. Bournazos, S. et al. Broadly neutralizing anti-HIV-1 antibodies require Fc effector functions for in vivo activity. Cell 158, 1243–1253 (2014).

37. Liu, Q. et al. Antibody-dependent-cellular-cytotoxicity-inducing antibodies significantly affect the post-exposure treatment of Ebola virus infection. Scientific reports 7, 45552 (2017).

38. Ao, Z., Ouyang, M.J., Olukitibi, T.A. & Yao, X. SARS-CoV-2 Delta spike protein enhances the viral fusogenicity and inflammatory cytokine production. iScience 25, 104759 (2022).

39. Yu, Y. et al. Antibody-dependent cellular cytotoxicity response to SARS-CoV-2 in COVID-19 patients. Signal Transduct Target Ther 6, 346 (2021).

40. Lee, W.S. et al. Decay of Fc-dependent antibody functions after mild to moderate COVID-19. Cell Rep Med 2, 100296 (2021).

41. Temming, A.R. et al. Cross-reactivity of mouse IgG subclasses to human Fc gamma receptors: Antibody deglycosylation only eliminates IgG2b binding. Molecular Immunology 127, 79–86 (2020).

42. Cao, J. et al. Development of an antibody-dependent cellular cytotoxicity reporter assay for measuring anti-Middle East Respiratory Syndrome antibody bioactivity. Scientific reports 10, 16615 (2020).

43. Simhadri, V.R. et al. A Human Anti-M2 Antibody Mediates Antibody-Dependent Cell-Mediated Cytotoxicity (ADCC) and Cytokine Secretion by Resting and Cytokine-Preactivated Natural Killer (NK) Cells. PloS one 10, e0124677 (2015).

44. Wang, R. et al. Therapeutic potential of a fully human monoclonal antibody against influenza A virus M2 protein. Antiviral research 80, 168–177 (2008).

45. El Bakkouri, K., et al. Universal vaccine based on ectodomain of matrix protein 2 of influenza A: Fc receptors and alveolar macrophages mediate protection. Journal of immunology (Baltimore, Md.: 1950) 186, 1022–1031 (2011).

46. Wellford, S.A. et al. Mucosal plasma cells are required to protect the upper airway and brain from infection. Immunity 55, 2118–2134.e2116 (2022).

47. Su, B. et al. Update on Fc-Mediated Antibody Functions Against HIV-1 Beyond Neutralization. Frontiers in Immunology 10 (2019).

48. Tauzin, A. et al. A single dose of the SARS-CoV-2 vaccine BNT162b2 elicits Fc-mediated antibody effector functions and T&#xa0;cell responses. Cell Host & Microbe 29, 1137–1150.e1136 (2021).

49. Ao, Z. et al. Incorporation of Ebola glycoprotein into HIV particles facilitates dendritic cell and macrophage targeting and enhances HIV-specific immune responses. PLoS ONE 14, e0216949 (2019).

50. Olukitibi, T., Ao, Z.-J., Mahmoudi, M., Kobinger, G. & X-j., Y. Dendritic cells/macrophages targeting feature of Ebola glycoprotein and its potential as immunological facilitator for antiviral vaccine approach.. Microorganisms 7, 402–425 (2019).

51. Olukitibi, T. et al. Development and characterization of influenza M2 ectodomain and/or HA stalk-based DC-targeting vaccines for different influenza infections.. Fronties in Microbiology **August** 08 (10.3389/fmicb.2022.937192) (2022).

52. Liu, W., Zou, P., Ding, J., Lu, Y. & Chen, Y.-H. Sequence comparison between the extracellular domain of M2 protein human and avian influenza A virus provides new information for bivalent influenza vaccine design. Microbes and Infection 7, 171–177 (2005).

53. Olukitibi, T.A. et al. Development and characterization of influenza M2 ectodomain and/or hemagglutinin stalk-based dendritic cell-targeting vaccines. Frontiers in Microbiology 13 (2022).

54. Ranadheera, C., Coombs, K.M. & Kobasa, D. Comprehending a Killer: The Akt/mTOR Signaling Pathways Are Temporally High-Jacked by the Highly Pathogenic 1918 Influenza Virus. EBioMedicine 32, 142–163 (2018).

55. Mlcochova, P. et al. SARS-CoV-2 B.1.617.2 Delta variant replication and immune evasion. Nature 599, 114–119 (2021).

56. Ao, Z., Fowke, K.R., Cohen, E.A. & Yao, X. Contribution of the C-terminal tri-lysine regions of human immunodeficiency virus type 1 integrase for efficient reverse transcription and viral DNA nuclear import. Retrovirology 2, 62 (2005).

57. Donofrio, G. et al. A Simplified SARS-CoV-2 Pseudovirus Neutralization Assay. Vaccines 9, 389 (2021).

58. Hu, J. et al. Development of cell-based pseudovirus entry assay to identify potential viral entry inhibitors and neutralizing antibodies against SARS-CoV-2. Genes & Diseases 7, 551–557 (2020).

59. Ouyang, M.J., Ao, Z., Olukitibi, T.A. & Yao, X.-J. Protocol to evaluate the inflammatory response in human macrophages induced by SARS-CoV-2 spike-pseudotyped VLPs. STAR Protocols 4, 102083 (2023).

60. Karakus, U., Crameri, M., Lanz, C. & Yángüez, E. in Influenza Virus: Methods and Protocols. (ed. Y. Yamauchi) 59–88 (Springer New York, New York, NY; 2018).

61. Reed, L.J. & Muench, H. A simple method of estimating fifty percent endpoints. American Journal of Epidemiology 27, 493–497 (1938).

